# Dendrites enable a robust mechanism for neuronal stimulus selectivity

**DOI:** 10.1101/023200

**Authors:** Romain D. Cazé, Sarah Jarvis, Amanda J. Foust, Simon R. Schultz

**Affiliations:** Center for Neurotechnology and Dept. of Bioengineering, Imperial College London, South Kensington, London SW7 2AZ, UK

**Keywords:** Dendrites, Stimulus selectivity, Synaptic integration

## Abstract

Hearing, vision, touch-underlying all of these senses is stimulus selectivity, a robust information processing operation in which cortical neurons respond more to some stimuli than to others. Previous models assume that these neurons receive the highest weighted input from an ensemble encoding the preferred stimulus, but dendrites enable other possibilities. Non-linear dendritic processing can produce stimulus selectivity based on the spatial distribution of synapses, even if the total preferred stimulus weight does not exceed that of non-preferred stimuli. Using a multi-subunit non-linear model, we demonstrate that stimulus selectivity can arise from the spatial distribution of synapses. We propose this as a general mechanism for information processing by neurons possessing dendritic trees. Moreover, we show that this implementation of stimulus selectivity increases the neuron's robustness to synaptic and dendritic failure. Importantly, our model can maintain stimulus selectivity for a larger range of synapses or dendrites loss than an equivalent linear model. We then use a layer 2/3 biophysical neuron model to show that our implementation is consistent with two recent experimental observations: (1) one can observe a mixture of selectivities in dendrites, that can differ from the somatic selectivity, and (2) hyperpolarization can broaden somatic tuning without affecting dendritic tuning. Our model predicts that an initially non-selective neuron can become selective when depolarized. In addition to motivating new experiments, the model's increased robustness to synapses and dendrites loss provides a starting point for fault-resistant neuromorphic chip development.

## 1 Introduction

The standard model of neuronal integration in neuroscience, which owes much to Hubel and Wiesel (Hubel and Wiesel, 1959), produces stimulus selectivity at the neuronal level by linearly integrating inputs within a single compartment. This model neglects the rich and in many cases spatially precise structure of the dendritic tree associated with many neuronal cell types throughout the brain (Stuart et al., 2016).

Several groups have recently presented data which is counter-intuitive given this standard model, as applied to orientation selectivity in the visual cortex. Firstly, this model integrates inputs with a narrow range of selectivity. In contrast, some experimental groups observed a mixture of selectivity (Smith et al., 2013; Jia et al., 2010). Specifically, Smith et al performed dual soma-dendrites recordings and they have demonstrated that somatic and dendritic tuning could differ (Smith et al., 2013). Moreover, Jia and colleagues have shown using calcium imaging that the tuning of dendritic hotspots could also differ from the somatic tuning (Jia et al., 2010). The first set of observations can be explained in a Hubel and Wiesel type model by using a higher number of synapses for preferred than non-preferred stimuli. Secondly, it was observed that hyperpolarization can significantly broaden somatic tuning, while dendritic tuning stays sharp (Jia et al., 2010). It is more difficult however to explain this second set of observations with a linear model. Why hyperpolarization does not also broaden dendritic tuning like it does for somatic tuning? Taken together these two sets of observations call for a new model and we propose here that these observations can be accounted for by the properties of dendrites.

Biophysical studies from the 80s and 90s demonstrated that a neuron can be sensitive to the spatial distribution of synaptic inputs because of its dendrites (Mel, 1993; Koch et al., 1982). Mel and colleagues have shown that a neuron could respond more intensely to clustered than dispersed inputs (Mel, 1993; Poirazi et al., 2003). Alternatively, Koch and colleagues have had demonstrated that under other conditions the opposite can also be true: a neuron can respond more to dispersed than clustered inputs (Koch et al., 1982). More recently it was also proposed that neuron can respond to a global stimulation (Poleg-Polsky, 2015) Our previous studies built on these biophysical findings and demonstrated that dendrites extend the computational capacity of a single neuron (Cazé et al., 2012, 2013). Recent experimental evidence has shown that excitatory synapses distribute non-randomly on dendrites and can form synaptic clusters (Takahashi et al., 2012; Druckmann et al., 2014; Kleindienst et al., 2011). We examine here whether we can employ the spatial distribution of excitatory synapses to implement stimulus selectivity. We show that such an implementation is more robust than a linear equivalent model and propose that it could better explain the recent experimental data.

## 2 Results

### Dendrites enable stimulus selectivity based on the spatial distribution of synapses

We show here how it is possible for a neuron to implement stimulus selectivity even if both the preferred and the non-preferred inputs make the same number of equal weight synaptic contacts.

We generate stimulus selectivity with a multi-subunit neuron model (see method) sensitive to the spatial distribution of synaptic inputs. Each subunit of this model nonlinearly transforms its synaptic input (Fig. 1A) similar to recent experimental observations (Polsky et al., 2004; Tamás et al., 2002; Abrahamsson et al., 2012; Cash and Yuste, 1998). A single large cluster of active synapses on a subunit generates a single dendritic spike whereas multiple smaller clusters of synapses (at least 40 synapses in this case) can generate multiple dendritic spikes. The neuron only fires when it receives multiple dendritic spikes. Because of this property, our model is more likely to fire an action potential when synaptic activation is distributed across multiple subunits rather than clustered onto a single subunit. We positioned the synapses to exploit this property and represent their distribution in Fig. 1B. The 100 synapses corresponding to the preferred stimulus distribute equally across two dendrites (50 onto subunit #1, 50 onto subunit #2 Fig. 2 top). In contrast, synapses from the non-preferred stimulus cluster primarily onto one dendrite (20 onto dendrite #1, 80 onto dendrite #2 Fig. 2 bottom). 20 synapses weakly drive dendrite #1, while 80 synapses saturate dendrite #2 (Fig. 2 bottom). Their summation of the dendritic integration at the soma falls short of the somatic spike threshold (red line Fig. 2). In contrast, the preferred stimulus’s synapses drive both dendrites such that their summation at the soma exceeds threshold. Thus, input ensembles with equal total synaptic weight can differentially affect the neuron’s spiking through their spatial distribution across dendrites.

**Figure 1:**
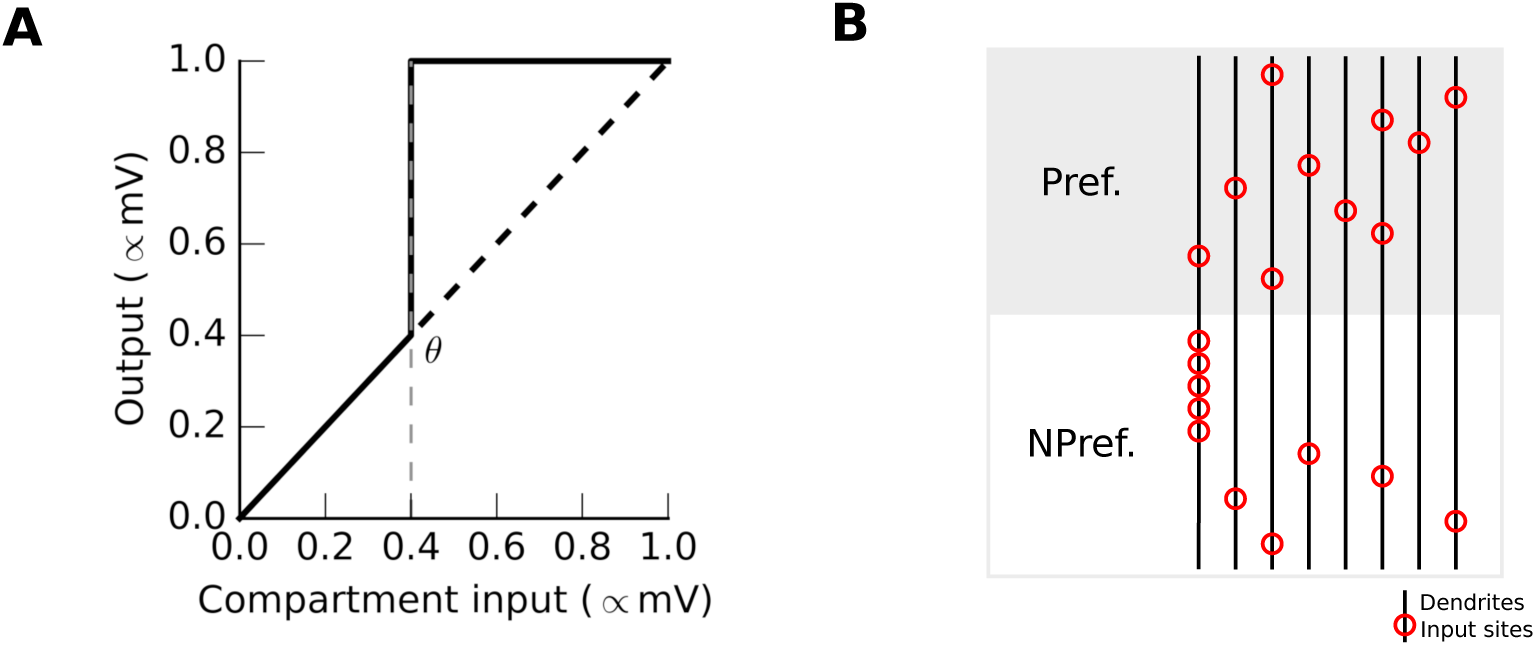
A non-linear model where activated synapses are distributed differently for preferred and non-preferred stimuli. (A) Local transfer function within a subunit; a non-linear jump occurs at the point *θ*, the non-linearity threshold. Input and output are normalized given their respective maxima. (B) The input sites (a red circle corresponds to 10 synapses) on dendrites (horizontal lines), a subunit corresponds to four branches. The preferred stimulus activates synapses distributed equally on both dendrites, whereas the non-preferred stimulus activates more synapses on the first subunit.

**Figure 2:**
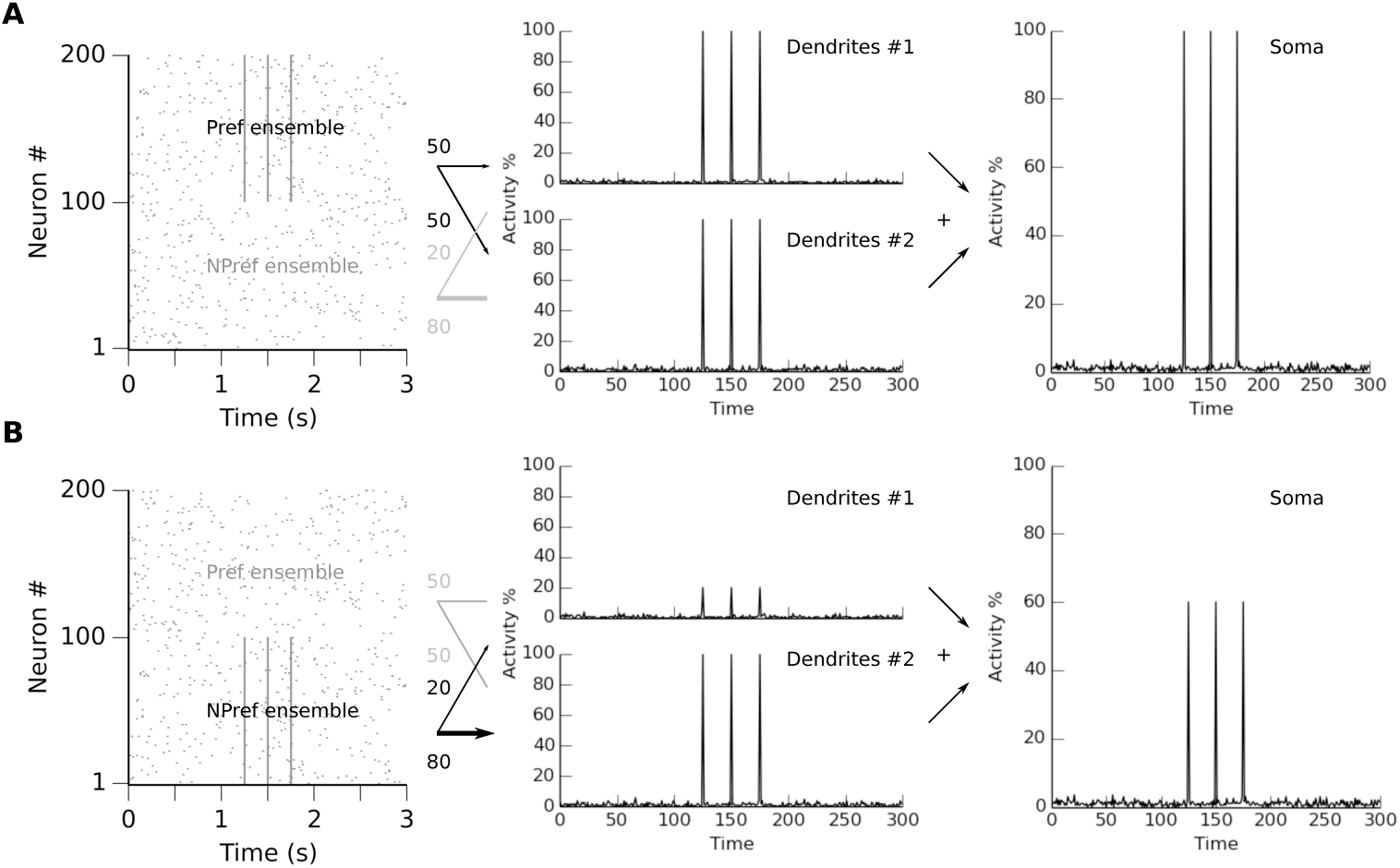
Implementing stimulus selectivity using the spatial distribution of synapses. *First column,* raster plot demonstrating the model’s input (top for the preferred, and bottom for non-preferred). Three synchronous events of a hundred spikes occur with stimuli that are preferred (A, neurons 101-200) and non-preferred (B, neurons 1-100). The number of synapses made by each ensemble are indicated above the arrows that show onto which of two dendrites they target. *Second column* result of dendritic integration within the two subunits. A dendritic spike occurs when more than 40 synapses activate on a subunit. *Last column* the somatic activity, an arithmetic sum of the dendritic sub-unit activity.

For a single preferred stimulus, the implementation could be directly done by connecting only the preferred stimulus’ synapses, all other synapses being irrelevant. We employ the spatial distribution of synapses in this case to demonstrate its use, but all synapses become relevant for slightly more complex and realistic cases. For instance, let us take three distinct stimuli: X, Y and Z. The neuron must respond to XY and XZ but not YZ. It has two non-linear subunits: X’s synapses targets the first subunit and Y/Z’s synapses target the second. X’s synapses saturates the first subunit and the second subunit saturates as soon as Y’s or Z’s synapses activate. The neuron will fire only if both subunits saturate. Consequently, the preferred stimuli XY and XZ will be more dispersed than YZ. In this case all synapses do matter and we use space to implement the function.

### Spatial distribution-based stimulus selectivity increases robustness to synaptic and dendritic failure

We compare in this section two types of models, linear and non-linear, with more than 5000 synapses of equal weight distributed non-deterministically. We benchmark how resilient they are to dendritic and synaptic losses. In both models, synapses distribute similarly onto seven dendritic subunits (see methods tab. 1), and the activity at the soma is the linear sum of all subunit activity. In the non-linear model, each subunit integrates synaptic inputs and 100 active synapses suffice to trigger a dendritic spike, saturating the subunit. In the linear model, however, synaptic integration results in the linear sum of all synaptic inputs. To compare these two models of synaptic integration we computed separability, the fraction of the model's instances capable of separating the preferred stimulus from all non-preferred stimuli. We generated a 1000 instances of the two models and count how many evoked activity in the soma larger for the preferred stimulus than all the non-preferred stimuli.

**Table 1.**
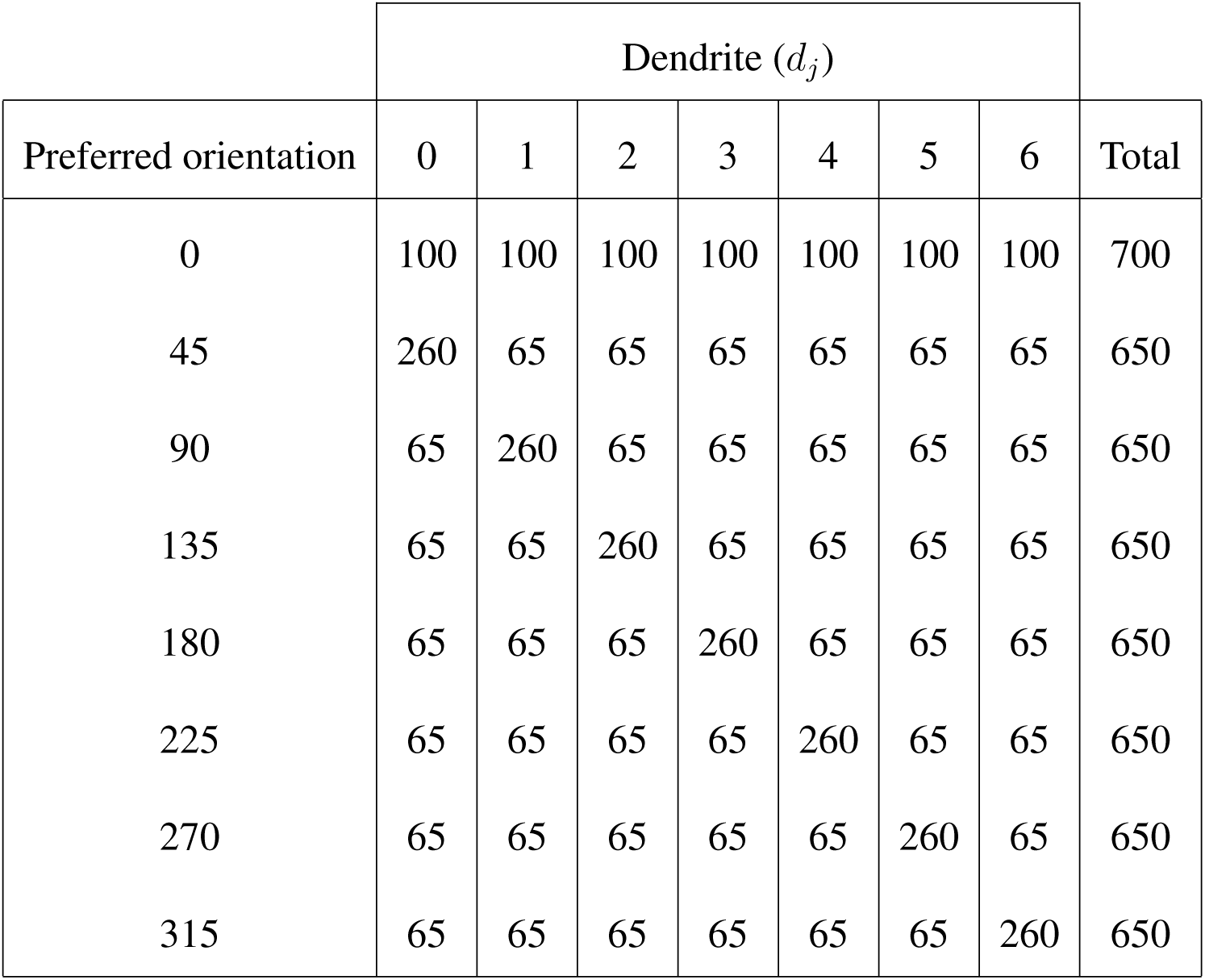
Synaptic distribution in our multi-subunit model. Mean number of synapses made by each presynaptic ensemble for each stimulus, for each postsynaptic dendrite.

We started by comparing the resilience of these models to the loss of synapses (Fig. 3). To maintain the same stimulus selectivity in the linear model, it is necessary that the ensemble coding for the preferred stimulus makes the strongest contact, e.g. makes the highest number of synaptic contacts or makes the synaptic contact with the highest weights. This prevents the linear model from selecting for a stimulus when another stimulus makes stronger contacts (proof in Methods). This is not the case for a neuron with non-linear dendrites. We can see that the non-linear neuron remains selective for the preferred stimulus even when the other stimuli have 150 more synapses (Fig. 3A, red). This property confers to the non-linear model robustness against synaptic failure (Fig. 3B). The non-linear model can separate both types of stimuli (Fig. 3C) and can maintain its function after 50% of its synapses fail (Fig. 3D). Given the same number of synapses the non-linear model will alway be more robust than the linear model.

**Figure 3:**
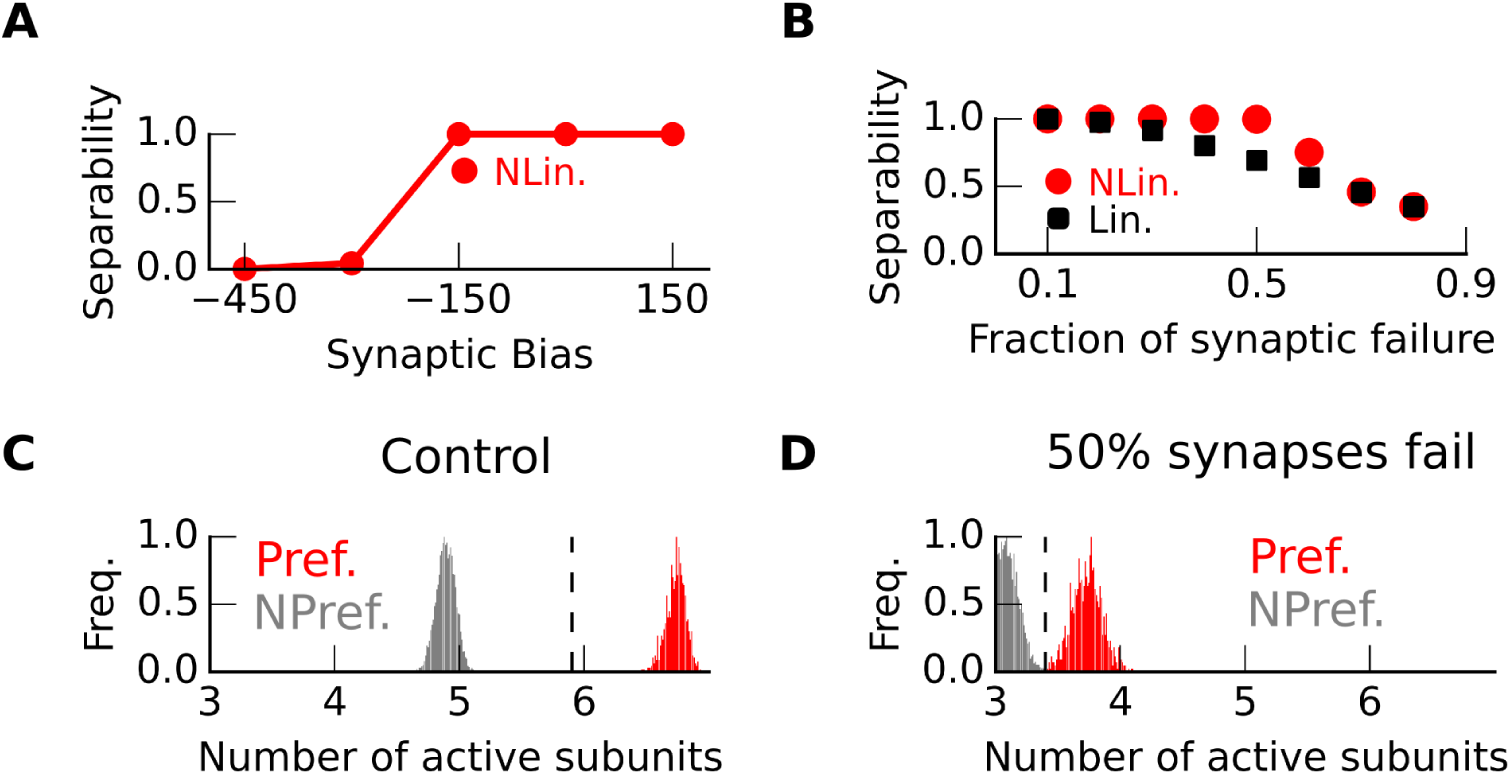
Stimulus selectivity achieved with spatial distribution of synapses increases the robustness to synaptic failure. (A-B) Separability, calculated as the fraction of model capable of separating preferred and non-preferred stimuli (over 1000 model following the synaptic distribution depicted in Table 1), for the non-linear (red circles) and linear (black square) models as a function of (A) the synaptic bias, which is the difference in the number of synapses between preferred and non-preferred ensemble; or (B) the synaptic failure which is the fraction of malfunctioning synapses. Here synaptic bias is zero. (C-D) Fraction of spiking/active subunits in a model with seven subunits (a subunit may not be fully active). Distribution for preferred (red) and non-preferred stimuli (gray) (C) In control condition, or (D) with 50% synapses failing.

If the number of synapses is the same for all ensembles, a linear neuron will never be able to separate preferred from non-preferred inputs. In the non linear model, however, stimulus selectivity arises through the clustering of synapses coming from the non-preferred ensembles (Fig. 4A red line). This way of implementing stimulus selectivity in the non-linear model fundamentally differs from the method used in a linear model.

**Figure 4:**
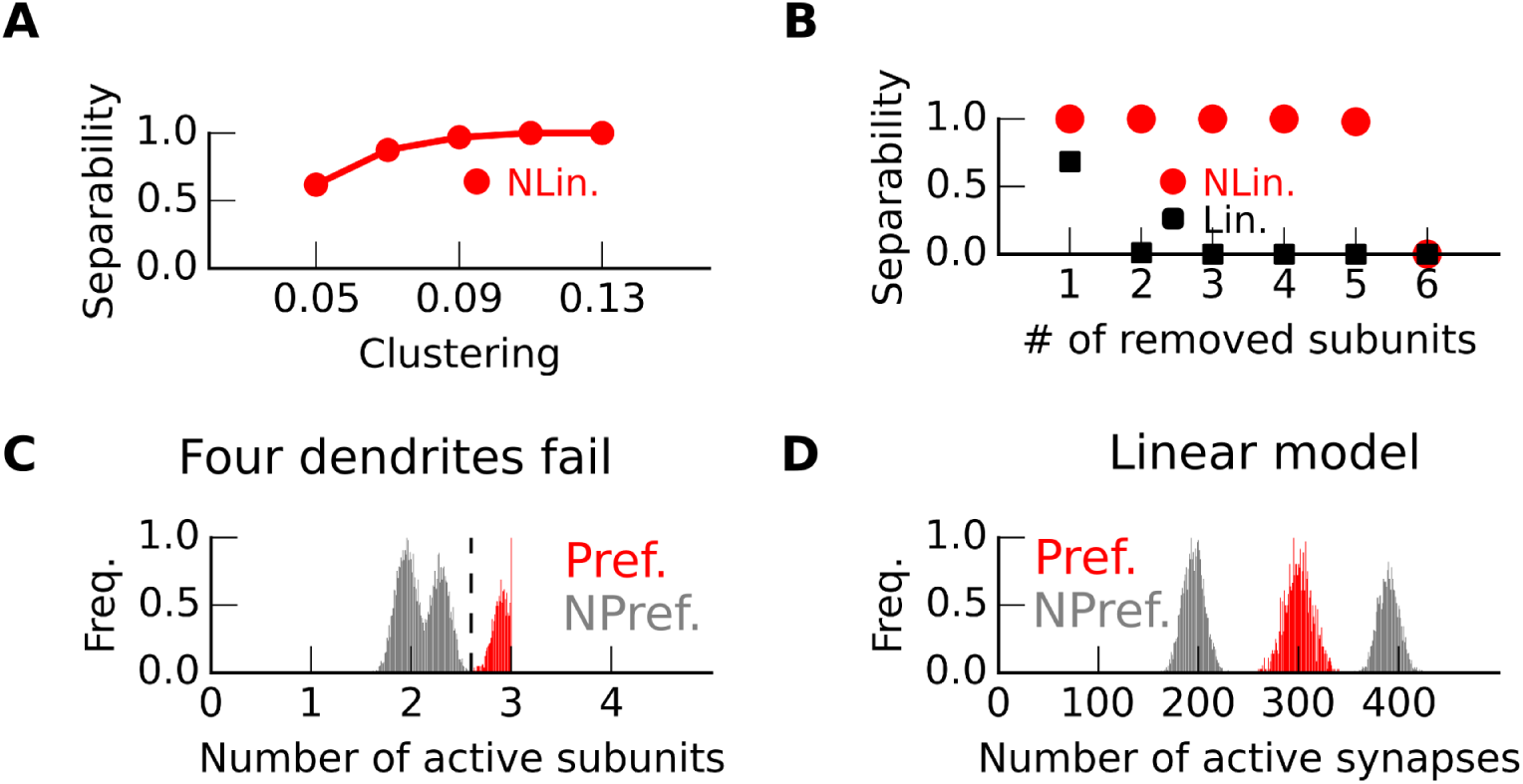
Stimulus selectivity achieved with spatial distribution of synapses increases the robustness to dendritic failure. (A-B) Separability, calculated as the fraction of models capable of separating preferred and non-preferred stimuli (*n* = 1000), for the non-linear (red circles) and linear (black square) model as a function of (A) the clustering bias, which is the number of synapses specifically set on a precise compartment; and (B) the number of removed subunits. (C-D) Distribution of dendritic activity for preferred (red) and non-preferred (gray) (C) in the non-linear models where dendritic activity closely relates to the number of maximally active compartment; and (D) in the linear model, where synaptic activity is the number of active synapses.

A non-linear model maintains its function when dendrites are disabled. This could occur when a dendrite is physically pruned from the neuron. Our multi-subunit nonlinear model maintains its function even if only 10 % of its synapses cluster on a single subunit as shown in Fig. 4A red). This considerably boosts the stability of the nonlinear model, which in this implementation can maintain functionality even with of loss of more than 50 % of compartments (Fig. 4B, black and Fig. 4C). In comparison, a linear model cannot make use of the spatial distribution of synapses. Therefore, if the synaptic bias is nil, it is impossible to differentiate preferred from non-preferred stimuli (Fig. 4A, black). The clustering bias (here 30 %) is detrimental for this type of model. It makes the linear model sensitive to the loss of even a single compartment (Fig. 4B black). Fig. 4D shows that, in this case, it is impossible for a linear model losing four dendrites to separate the preferred from non-preferred stimuli.

In summary, we compared two multi-subunit models: a linear and a non-linear model. We used simulations to demonstrate that the non-linear model is much more robust than its linear equivalent. We have shown that non-linear dendrites offer a new dimension of robustness. Our non-linear multi-subunit model can lose 50% (more than 2600) of its synapses or more than 50% of its dendrites (more than 4) while maintaining its function.

### The biophysical model replicates the mixture of dendritic tunings

We have shown how a multi-subunit non-linear model can robustly implement stimulus selectivity. The following section demonstrates that these results carry over to a biophysical model capturing rich temporal dynamics and interactions between compartments. This biophysical model fits recent experimental observations (Jia et al., 2010).

We constructed a stimulus selective neuron (Fig. 5A) that replicates the experimental data. Both the data and our model can display a variety of dendritic tunings (Fig. 5A-B). To replicate the experimental observations, we used 8 ensembles of AMPA/NMDA-type synapses distributed each on 7 locations. Synapses from the preferred stimulus ensemble scatter across all branches, whereas synapses from the non-preferred ensembles cluster, each onto a particular dendrite. We placed these synapses on a Layer 2/3 neuron reconstruction (Jia et al., 2010).

**Figure 5:**
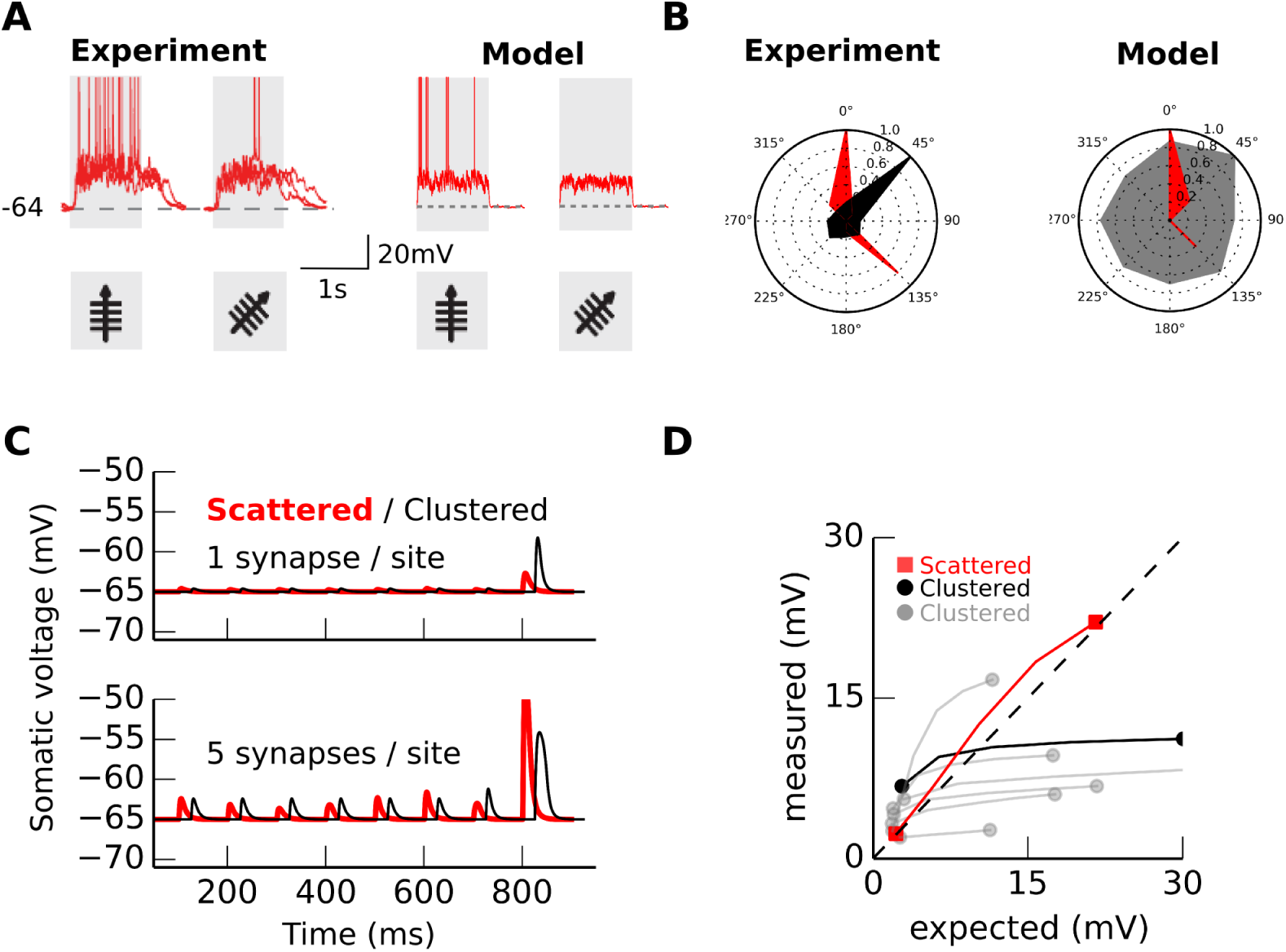
Stimulus selectivity using our implementation displays a mixture of dendritic selectivities. (A) Somatic voltage for two stimuli (0/45 degrees during shaded period). (B) Spike count (red) and experimentally determined integral of the calcium response in dendrites (black) vs integral of the voltage response in dendrites (one example, gray). Note the similarity of the somatic tuning but not of the dendritic tuning. The experimental’s sharp tuning is due to hyperpolarization. (C) Somatic depolarization when one(top)/five(bottom) synapse(s) activate in one of the input sites from 0 to 800 ms; or when one/five occurs at all the 7 sites (800 to 1000ms). (D) Expected (arithmetic sum) versus measured depolarization in the 8 sets (red:scattered on 7 branches, black/grey:clustered on a branch) of 7 input sites. Black and red marks correspond to experiments carried out in (C). The four dots (two squares and two circles) correspond to the four pairs (scattered/clustered case).

The activation of a synapse results in a somatic depolarization of 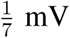, independent of its location (Fig. 5C), as has been observed in another cell type (Smith et al., 2003). We enforced this “dendritic democracy” (Häusser, 2001) by scaling synaptic conductances depending on their distance to the soma. Consequently, all synapses produce the same depolarization at the soma and each ensemble makes the same number of synapses. All ensemble may therefore produce the same depolarization at the soma, this is not what is happening because of non-linear interaction between synapses.

Interestingly, synapses interact in two distinct ways, dependent on their location. For synapses clustered on a branch (Fig. 5C black line), seven active synapses, one per location interact supra-linearly and they produce a depolarization superior to one mV because they generates an NMDA spike (Nevian et al., 2007), but 35 synapses on a branch (five per location) interact sub-linearly due to reduced driving force at synapses (Koch et al., 1982; Tran-van Minh et al., 2015). Even if the depolarization at the soma is weak, locally, the membrane voltage within a branch reaches the equilibrium voltage (0 mV) because of the branch small diameter 1 *μ*m (see Movie S1). In contrast, for 35 synapses distributed across the seven branches, scattered stimulation depolarizes the soma more than the same number of clustered synapses because their activation generates multiple NMDA spikes (Fig. 5C, red line) as has been observed experimentally *in vivo* (Jia et al., 2014). These observations are summarized in an expected/measured plot (Fig. 5D) and show the biophysical model's sensitivity to the synaptic spatial distribution.

This sensitivity enables the generation of stimulus selectivity in our model. If the population coding for the preferred stimulus makes functional synapses on all primary dendrites, whereas non-preferred stimuli cluster on a single branches, then the distributed synaptic arrangement produces multiple NMDA spikes that reach the soma in parallel, as observed in *vitro* (Larkum et al., 2007) and in *vivo* (Hill et al., 2013; Palmer et al., 2014) (Fig. 5A). Both scenarios are illustrated in animations provided as supplementary material (see movies S1 and S2). Because the preferred stimulus produces multiple NMDA spikes it generates the highest synaptic depolarization.

In a single compartment model, the highest weighted stimulus always ”wins”, rendering synaptic spatial distribution irrelevant. Conversely, our multi-compartment biophysical model uses exclusively the spatial distribution of synapses to implement stimulus selectivity, a configuration that could explain, in contrast with single compartment models, how calcium hotspots in dendrites display mixed stimulus tuning (Jia et al., 2010). Note that our model does not exclude an average dendritic tuning similar to the somatic tuning, however it can explain the cases where the average dendritic tuning differs from somatic tuning. Although here we implemented spiking without dendritic backpropagation, our previous Hodgkin and Huxley-based implementation that allowed backpropagation exhibited the same result (data not shown).

### Hyperpolarization broadens somatic but sharpens dendritic tuning in our model

We injected current at the soma in our biophysical model to pull down the membrane potential from −65mV to −70mV, as in Jia et al.’s experiment (Jia et al., 2010). Because of hyperpolarization, the neuron stops firing action potentials, and the somatic tuning of the membrane voltage becomes broader than the tuning of spikes. This might be mainly due to the non-linearity induced by somatic spiking in the control condition. The dendritic tuning, however, is sharp even under hyperpolarization.

When we decrease the resting membrane voltage to −70mV, the number of synapses necessary to trigger a membrane non-linearity increases, and only the dendritic preferred stimulus provokes this non-linearity. Fig. 6A shows that only the 45 degree stimulus triggers this non-linearity, and consequently dendritic selectivity sharpens (Fig. 6B). Furthermore, this could be reinforced by the non-linearity of the calcium sensor. Conversely, the somatic depolarization difference between scattered and clustered synapses decreases, when we hyperpolarize the neuron (Fig. 6C-D), and somatic selectivity broadens.

**Figure 6.**
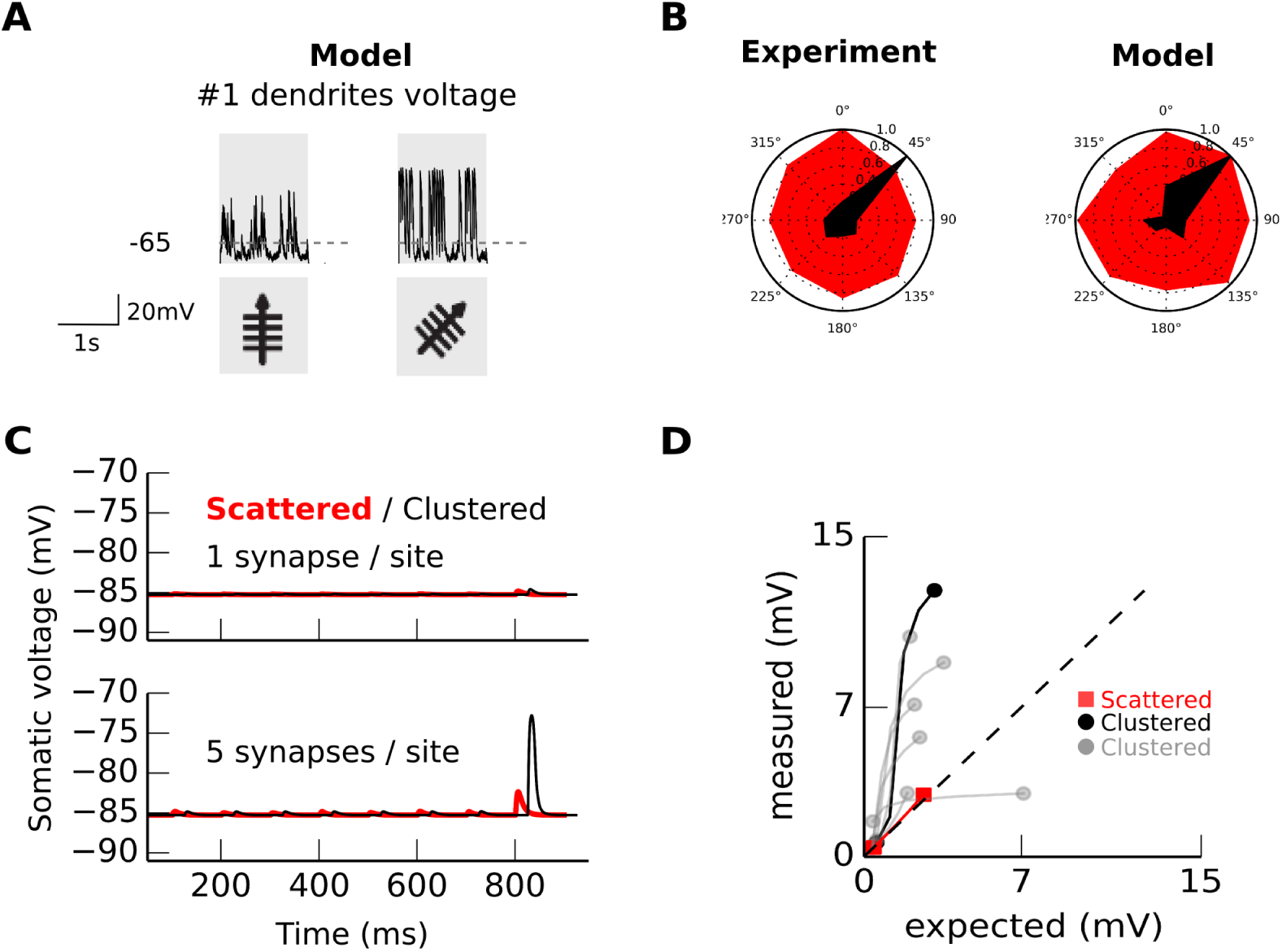
Hyperpolarization broadens somatic tuning whereas it sharpens dendritic tuning in our model. (A) Local membrane potential in the first dendritic branch for two stimuli: the soma’s preferred stimulus and for the dendrite’s preferred stimulus. (B) Integral of dendritic calcium signal (black) and of the somatic membrane voltage (red) (experimental data replotted from (Jia et al., 2010)). In our model we compute the integral of the dendritic membrane voltage. (C) Somatic depolarization when one(top) or five(bottom) synapses activate in one of the seven input site from 0 to 800 ms; or when one/five activate at all the seven sites simultaneously (800 to 1000ms). (D). Expected (arithmetic sum) versus measured depolarization in the simulation 2 sets of 7 locations.

The model's sensitivity to the spatial distribution of synapses predicts the effect of hyperpolarization on dendritic tuning. The broadening of the somatic tuning can intuitively be explained by hyperpolarization. Intuitively, somatic spiking non-linearly sharpens somatic tuning in the control condition. The sharp dendritic tuning however is much less intuitive, and the sharpening of dendritic tuning during hyperpolarization is another important prediction of our modeling work. This could be tested by using micro-injection of TTX, or a similar approach, instead of hyperpolarization to block back-propagated action potentials.

## 3 Discussion

We implemented stimulus selectivity in a multi-subunit non-linear model (Fig. 2 & Fig. 1). This implementation of stimulus selectivity is more robust than the linear one (Fig. 3 & Fig. 4). Because it is possible in our model that the preferred stimulus can be less strongly connected than non-preferred stimuli, our implementation is resilient to synapse or dendrites loss.

In our model, the preferred stimuli generate multiple clusters and elicit multiple NMDA spikes whereas the non-preferred inputs elicit a single NMDA spike. We implemented NMDA spikes for biological relevance, but a saturating dendritic non-linearity would have also sufficed. The dendritic non-linearity is the main parameter responsible for the enhanced robustness. The amount of added robustness depends on the model parameters, e.g. the number of dendrites (Fig. /reffig4). Future work can study in detail the influence of parameters such as somatic threshold and subunit number.

Our non-linear implementation can coexist with the classic implementation based upon synaptic strength (Fig. 5A), providing an additional channel for neuronal information processing. As it has been observed, the ensemble encoding the preferred stimulus does make the strongest contact, as suggested by Cossel et al (Cossell et al., 2015) and observed by Chen et al (Chen et al., 2013). We provide here a new layer of robustness through the spatial distribution of synapses.

Locally non-linear integration could have made our model cluster sensitive. A neuron with the latter type of sensitivity might possess the same robustness to synaptic failure than our neuron model, but not to dendritic failure. Instead the neuron model used here is scatter sensitive: it responds more to scattered (widely distributed) than clustered synaptic activity. This behavior has previously been described (Koch et al., 1982; Mel, 1993), but has never been proposed as a mechanism underlying stimulus selectivity. Scatter sensitivity, contrary to cluster sensitivity, only requires saturating non-linearities (Cazé et al., 2012).

Additional experimental work could confirm the functional role of scatter sensitivity and show that stimulus selectivity is resilient to dendrites removal. For instance, targeted laser dendrotomy (Go et al., 2016).

Importantly, the average dendritic tuning and somatic tuning can, different to the results presented here, be identical. Let’s take a case where each branch does not have the same weight and produces the same depolarization on the soma. One could imagine a situation where branches with a small weight have the same tuning as the soma. If these branches are the most numerous, then the average dendritic tuning will correspond the somatic tuning.

Although we have focused, to ease comparison with experimental studies, on the tuning of a neuron to sensory stimuli, all neural computations can be described in terms of stimulus selectivity. Boolean functions can both describe a neural computation and stimulus selectivity. In the latter case, we can describe stimuli as words of 0s and 1s. In the former case, we can describe all neural computations as Boolean functions if we binarize activity. Therefore our implementation based on the spatial distribution of synapses can be used for general neural computation. To transpose a synaptic strength based implementation of a computation, it suffices to turn inputs with the strongest synapses activated into inputs with the most dispersed synapses activated.

The biophysical model reinforces our work that the insights gained from the multisubunit model are physiologically relevant; together, they yield three predictions. Firstly, we predict that hyperpolarization not only broadens somatic tuning (Jia et al., 2010; Lavzin et al., 2012) but it also sharpens dendritic tuning (Fig. 6). Secondly, our model predicts that a neuron may recover its tuning after losing a large fraction of either its synapses or dendrites, due to the robustness provided by spatial synaptic distribution based information processing. Thirdly, we predict that a cortical neuron with no apparent stimulus tuning can acquire stimulus selectivity when depolarized, similar to what can be observed in place cells (Lee et al., 2012).

Our implementation using nonlinear dendritic integration, that can be learned using an unsupervised learning algorithm (Cazé et al., 2016), may inform the design of neuromorphic chips, as it suggests that the use of dendrites-even passive-can extend the robustness of the circuit. While we have demonstrated these capabilities in the context of a model neuron’s selectivity to a visual stimulus, the mechanism we have proposed is general, and potentially reflects a canonical computational principle for neuronal information processing. Whether or not the mechanism proposed here turns out to underlie or assist in selectivity to the orientation of visual stimuli, it may be examining further in the study of a range of elementary information processing operations involving neurons with rich dendritic trees.

## 4 Methods

### Multi-subunit model

Our neuron model is made of subunits *D_j_*, each sums their synaptic inputs 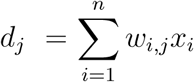 where *x_i_* and *w_i,j_* are equal to 0 or 1:

- In the linear case *D_j_*(*d_j_*) = *d_j_*
- In the non-linear case 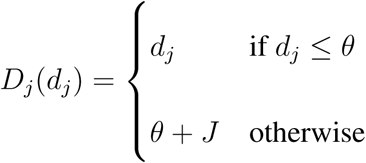. See Fig. 1A

This results in a somatic activity equals to 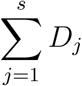

For the elementary model used in the first part *θ* = 40 and *J* = 60 and in the model with seven subunits and more synaptic inputs *θ* = 100 and *J* = 0 to show that a saturating non-linearity suffices to exploit the spatial distribution of synapses.

In this latter model the seven subunits receive input coming from eight presynaptic neuronal ensembles corresponding to eight different stimulus orientations. The mean number of synaptic contacts for each ensemble-dendrite pair is described in Table 1. The preferred stimulus (0 degrees) activates 700 synapses following a random uniform distribution across all seven dendrites. In contrast, non-preferred stimuli activate 650 connections each, including a bias such that 40% of input from each orientation preferentially target one of the dendrites and the remaining 60% being uniformly distributed among the remaining six dendrites following a uniform distribution.

Two recent papers describe how such a spatial distribution of synapses could be learned (Legenstein and Maass, 2011; Wu and Mel, 2009). Please note that we also proposed a learning algorithm presented in a self-archived manuscript currently under review (Cazé et al., 2016).

### A necessary condition for the linear model

The highest weight needs to be from the preferred stimulus in a linear model. To prove that let us consider the simplest scenario where two presynaptic neurons each synapse onto a postsynaptic neuron. We arrange it so that one input codes for the preferred stimulus while the other for a non-preferred stimulus, and *W*_pref_ and *W*_nonpref_ is the amplitude of their resulting depolarization on the postsynaptic neuron. Here, stimulus selectivity is possible only if *W*_pref_ ≥ Θ and *W*_nonpref_ < Θ, which is equivalent to *W_pref_* > Θ > *W*_nonpref_. This condition can be generalized for any number of presynaptic neurons, and implies in the linear neuron model when constrained to positive values of *W* that stimulus selectivity is only possible when the preferred stimulus has the highest weight.

### Biophysical model

For detailed modeling, we used a reconstructed morphology of a neuron from Layer 2/3 of visual cortex in mouse (Jia et al., 2010). The capacitance of the model is *C* = 1*μF/cm*^2^. The axial resistance in each section was *R_a_ =* 100Ω.*cm* to match the one observed in pyramidal neurons (Stuart and Spruston, 1998; Oswald and Reyes, 2008), and passive elements were included (*g_l_* = 0.0003Ω^−1^, *e_l_* = −65 mV). To hyperpolarize the neuron to −85*mV* we injected in the soma 0.2*nA* which gives an input resistance of 100MΩ matching what is observed in pyramidal neurons. Spiking was implemented using an integrate-and fire mechanism with a hard threshold of −40mV, which has been shown to provide an accurate depiction of spike initiation behaviour (Brette, 2015), whereupon we set the voltage to 20 mV in the following timestep, before resetting to 65 mV. We have previously used a Hodgkin Huxley for the generation of action potential and we obtained the same result (data not shown). The model was implemented using NEURON with a Python wrapper (Hines et al., 2009), with the time resolution set to 0.1ms.

### Synaptic inputs to the biophysical model

We used 280 synapses divided into 8 groups of 35 synapses, corresponding to 8 different stimuli (orientations). Each had a background activity of 1 Hz which increased to 10Hz during the presentation of the stimulus. As experimental evidence suggests that stimulus information is coded not only by an increase in firing rate but also in correlation (Bruno and Sakmann, 2006; DeCharms and Merzenich, 1996), synapses synchronously coactivate 20 times to encode the presence of a stimulus (preferred or otherwise). This raises the firing rate of this group to 30 Hz. The specific set of synchronous synapses activated depends on the stimulus identity, e.g. synapses 1-35 synchronously activate for the preferred stimulus, synapses 36-70 activate for the non-preferred stimulus #1, etc.

### Conductance based NMDA-type synapses

NMDA-like inputs were included by modeling voltage-dependent, conductance-based synapses that generated a postsynaptic currents *i_s_* = *g*(*t*)*g*_mg_(*v*) × (*v*(*t*) − *e_s_*), with reversal potential *e_s_* =0 mV. For *g*(*t*), we used a two scheme kinetic scheme with rise and decay time constants *τ_1_* = 0.1ms and *τ_2_* = 10ms (Destexhe et al., 1994). The voltage-dependent conductance *g_mg_ (v)* was determined assuming [Mg^2+^] = 1mM. The equation for the *Mg* block was 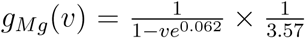 following (Jahr and Stevens, 1990). Files we used to run the binary and biophysical simulations are available on the github repository of the corresponding author (github.com/rcaze).

## Supporting Information

### S1 Video

**Neuron response to clustered synaptic inputs.** L2/3 neuron reconstruction from the visual cortex. The large circle is the soma and black dots are input sites. Depolarisation of a section is color-coded (black:low, yellow:high).

### S2 Video

**Neuron response to scattered synaptic inputs.** L2/3 neuron reconstruction from the visual cortex. The large circle is the soma and black dots are input sites. Depolarisation of a section is color-coded (black:low, yellow:high).

## Acknowledgments

The authors thank Dr Hongbo Jia for providing the Layer 2/3 neuron reconstruction used for the model presented in this paper, and Dr. Andrew Gallimore, Dr. Mark Humphries, Dr. Boris Gutkin, Dr. Fleur Zeldenrust and Dr Matthijs Van Der Meer for their comments on the draft manuscript. This work was supported by EU FP7 Marie Curie Initial Training Network 289146. SJ is supported by EU FP7 Marie Curie fellowship (PIEF-GA-2013-628086), and SRS by a Royal Society Industry Fellowship.

